# *Clostridioides difficile* phosphoproteomics shows an expansion of phosphorylated proteins in stationary growth phase

**DOI:** 10.1101/2021.11.11.468335

**Authors:** Wiep Klaas Smits, Y. Mohammed, Arnoud de Ru, Valentina Cordó, Annemieke Friggen, Peter A. van Veelen, Paul J. Hensbergen

## Abstract

Phosphorylation is a post-translational modification that can affect both house-keeping functions and virulence characteristics in bacterial pathogens. In the Gram-positive enteropathogen *Clostridioides difficile* the extent and nature of phosphorylation events is poorly characterized, though a protein-kinase mutant strain demonstrates pleiotropic phenotypes. Here, we used an immobilized metal affinity chromatography strategy to characterize serine, threonine and tyrosine phosphorylation in *C. difficile*. We find limited protein phosphorylation in the exponential growth phase but a sharp increase in the number of phosphopeptides after the onset of stationary growth phase. Among the overall more than 1500 phosphosites, our approach identifies expected targets and phosphorylation sites, including the protein kinase PrkC, the anti-sigma-F factor antagonist (SpoIIAA), the anti-sigma-B factor antagonist (RsbV) and HPr kinase/phosphorylase (HprK). Analysis of high-confidence phosphosites shows that phosphorylation on serine residues is most common, followed by threonine and tyrosine phosphorylation. This work forms the basis for a further investigation into the contributions of individual kinases to the overall phosphoproteome of *C. difficile* and the role of phosphorylation in *C. difficile* physiology and pathogenesis.

**Importance:** In this manuscript, we present a comprehensive analysis of protein phosphorylation in the Gram-positive enteropathogen *Clostridioides difficile*. To date, only limited evidence on the role of phosphorylation in regulation in this organism has been published; the current study is expected to form the basis for research on this post-translational modification in *C. difficile*.

## INTRODUCTION

Proteins in organisms from all domains of life can be functionally altered by post-translational modifications (PTMs) that are often species-dependent (1). PTMs in bacteria generally occur at much lower levels than in eukaryotes (2). PTMs serve to regulate activity in response to environmental conditions, as they process generally faster than transcription-translation (2), but can also be used to tune phenotypic diversity (3). Amongst the high variety of PTMs (1), protein phosphorylation is one of the best-studied examples, partially attributable to major developments in phosphoproteomics.

To date, phosphorylation has been identified on the side chains of different amino acids: serine (Ser), threonine (Thr), tyrosine (Tyr), histidine (His), arginine (Arg), lysine (Lys), aspartate (Asp) and cysteine (Cys). The chemistry involved in the phosphorylation of different amino acid is different. Phosphorylation on Ser/Thr/Tyr results in more stable phosphoester bonds, whereas His/Lys/Arg phosphorylation results in thermodynamically unstable phosphoamidates (1). Phosphorylation on Asp leads to a mixed phosphoacylanhydride and modification of cysteine (Cys), finally, leads to a phosphothiolate (1). Ser/Thr/Tyr phosphorylation appears to be the most common in bacteria (4), and may relate to their thermodynamic stability.

In canonical protein phosphorylation, a phosphate group is transferred to a protein through the action of a kinase, and can be removed by a phosphatase; sometimes the kinase and phosphatase functions are encoded by the same bifunctional enzyme (1). Oftentimes, the response of bacteria to environmental stimuli is dependent on so called two component systems, that consist of a membrane-associated two-component sensor histidine kinase (HK) and a cytosolic response regulator (RR). When triggered, autophosphorylation of the HK on a histidine residue results in the transfer of the phosphate group to an aspartate residue on the response regulator. Subsequently, the phosphorylated RR can bind to DNA to regulate transcription of target genes (1). Other His-phosphorylated proteins include the phosphoenolpyruvate-dependent sugar phosphotransferase system (PTS) proteins, with the exception of the EIIB component that can be phosphorylated on Cys residues (5). As the name suggests, the donor for the phosphorylation of PTS systems is not a kinase, but phosphoenol pyruvate (PEP). The PTS phosphorylation cascade includes an unusual bifunctional kinase/phosphatase, called HPr, that in its S-phosphorylated form can interact with the transcription factor CcpA to regulate metabolic activity (5).

Phosphorylation on Ser/Thr is mediated by eukaryotic-type Ser/Thr Kinases (eSTK), also known as Hanks-type kinases (6). Interestingly there is some evidence that eSTKs may outnumber two component systems (7). Phosphorylation on Tyr in bacteria is mediated by a unique family of proteins known as BY kinases (2, 8, 9).

The exploration of bacterial phosphorylation has been greatly stimulated by the development of mass spectrometry-based phosphoproteomics techniques that provide a snapshot of the site-resolved phosphorylation state of proteins under given conditions (10, 11). These techniques rely on strategies to enrich for phosphorylated peptides, through for instance anti-phospho antibodies, strong cation exchange chromatography, chemical modifications or immobilized metal affinity chromatography (IMAC) (2, 4). Overall, the picture that emerges from these experiments is that both the number of phosphorylation sites and the relative abundance of different types of phosphorylation differs between organisms (2, 4, 11, 12). Additionally, it was found that one protein can be dynamically phosphorylated on multiple different sites (4, 13).

In bacteria, phosphorylation has been implicated in diverse processes such as cell cycle regulation, cellular differentiation, morphogenesis, and metabolism, persistence and virulence (1). Whereas phosphoproteomics initially focused on model organisms such as the Gram-negative *Escherichia coli* (14) and Gram-positive *Bacillus subtilis* (15), phosphoproteomes have been determined for a large number of pathogens as well (16, 17). Though some of the effects on pathogenicity can be related back to pleiotropic effects on cellular integrity, for instance, phosphorylation is also directly implicated in virulence by modulating virulence gene expression or factors involved in host establishment, as well as affecting the host immune system (10, 16). Finally, phosphorylation has also been found to affect antimicrobial resistance in multiple pathogens (16, 18). Together, these findings suggest that targeting phosphorylation might be a viable strategy for novel antimicrobial or anti-virulence strategies (16, 18).

Despite the wealth of phosphoproteomic studies (16, 17), phosphorylation in Clostridia is largely unexplored. A single study has been performed in *Clostridium acetobutylicum*, a solventogenic species with biotechnological relevance, that identified a total of 61 phosphorylated proteins (19). To our knowledge, no studies have been carried out in pathogenic species such as *Clostridioides difficile*.

*C. difficile* is a major cause of healthcare-associated diarrhea, and can cause a potentially fatal disease (*C. difficile* infection, or CDI) as a result of the production of toxins that affect the integrity of the colon epithelium (20-22). The development of CDI is clearly linked to reduced diversity of the microbiota of the host and consequently restoring this diversity for instance by fecal microbiota transplantations has proven to be an effective treatment (23, 24). Transmission and persistence of *C. difficile* is dependent on its ability to form endospores, as in a mutant unable to form spores these processes are severely inhibited (25). All the processes above are subject to complex regulation and - based on homology with the Gram-positive model organism *B. subtilis* – can be expected to involve phosphorylation of key proteins (26-30). However, important differences in these regulatory pathways exist between *B. subtilis* and *C. difficile* (31, 32). Importantly, genetic investigations have revealed that the eSTK protein PrkC of *C. difficile* is reported to affect cell wall homeostasis and resistance to antimicrobials, though the molecular mechanism through which this occurs have not been elucidated (33). In contrast, it was recently shown that PrkC directly affects cell division through phosphorylation of the peptidoglycan hydrolase CwlA (34). To date, this is the only reported substrate of the PrkC kinase.

Here, we report the first phosphoproteomic analysis of *C. difficile* based on an IMAC-based enrichment strategy. We show that phosphorylation is growth phase-dependent and abundant in stationary growth phase, and that several aspects of phosphorylation-dependent regulation appear to be conserved. Our results contribute to our understanding of phosphorylation in pathogens and pave the way for functional dissection of the role of kinases and phosphatases in *C. difficile*.

## MATERIALS AND METHODS

### Chemicals

Unless noted otherwise, chemicals were obtained from Sigma Aldrich Chemie.

### Cell culture

*C. difficile* 630Δ*erm* cells (35, 36) were grown at 37 °C in brain-heart infusion medium (BHI; Oxoid) supplemented with yeast extract (YE) in a Don Whitley VA-1000 workstation (10% CO_2_, 10% H_2_ and 80% N_2_ atmosphere). An initial pre-culture was grown until an optical density at 600 nm (OD_600nm_) of 0.36. Three independent cultures were started by adding 31 mL of the pre-culture to 400 mL fresh BHI-YE for each new culture. Samples were taken at mid-exponential (3.25h post inoculation), early stationary (5.25h post inoculation) and late stationary phase (24h post inoculation) (**Supplemental Figure 1**). To get a roughly equivalent amount of cells, 100 mL was taken at the mid-exponential time point (OD_600nm_ 0.96), while 50 mL was taken at the two later time points (OD_600nm_ ~ 2).

JY cells were grown at 37 °C in Iscove’s modified Dulbecco’s medium, supplemented with 10% heat-inactivated fetal bovine serum and L-glutamine. Both eukaryotic and bacterial cells were harvested by centrifugation (3184 x g, 10 min), and washed three times with 25 mL of phosphate buffered saline (PBS; Fresenius Kabi Nederland BV). Cell pellets were subsequently stored at -20 °C until further use.

### Sample preparation

Cell pellets were re-suspended in 3 mL lysis buffer (8 M urea/50 mM Tris-HCl pH 7.4, 1 mM orthovanadate, 5 mM Tris(2-carboxyethyl)phosphine (TCEP), 30 mM chloroacetamide (CAA), phosphoSTOP phosphatase inhibitor (Thermo Fischer Scientific), cOmplete™ mini EDTA free protease inhibitor, 1 mM MgCl_2_) and incubated for 20 min at room temperature. Cells were lysed by sonication (Soniprep 150 ultrasonic disintegrator (MSE), 5x 30 s, amplitude 12 microns). In between sonication steps samples were cooled on ice for 30 s. Next, samples were centrifuged for 15 min at 7200 x g at 4 °C. The urea concentration was adjusted to 6 M by the addition of 1 mL of 50 mM Tris-HCL pH 7.4, after which 1 μL Benzonase (250 U/ μL) was added and samples were incubated for two hrs at room temperature.

Proteins were precipitated by first adding 16 mL of methanol (Actu-All Chemicals) and mixing, followed by 4 mL of chloroform (Merck Millipore) and mixing. After the addition of twelve mL of Milli-Q water (obtained from a Elga Pure Lab Chorus 1 Machine), the samples were mixed by vortexing. Samples were then centrifuged for 15 min at 11000 g, followed by another 5 min at 11000 x g (this resulted in better separation of the two phases than one round of 20 min of centrifugation). The protein precipitates at the interphase were collected and washed twice with 2 mL methanol, followed by 5 min centrifugation at 11000 x g. The protein pellets were air-dried and re-suspended in 5 mL 25 mM NH_4_HCO_3_ pH 8.4.

Trypsin was added at a ratio of 1:25 (w/w) and overnight digestions were performed at 37 °C. The next day, samples were centrifuged for 10 min at 11000 x g and the supernatants were collected. Desalting of the samples was performed using Oasis HLB 1 cc Vac cartidges (Waters). Briefly, the cartridge was washed once with 1 mL acetonitrile (Actu-All Chemicals)/H_2_O 90/10 (v/v), equilibrated with Milli-Q H_2_O/acetonitrile/formic acid (Fluka Analytical) (95/3/0.1 v/v/v) (solution A, 2x 1 mL). Following sample loading, the column was washed with solution A (3x 1 mL) after which peptides were eluted with acetonitrile/water/formic acid (30/70/0.1 v/v/v). Tryptic peptides were lyophilized (Salm en Kipp Christ RVC 2-18 CD plus) and stored at -20 °C until use.

### Phosphopeptide enrichment

A one-step phosphopeptide IMAC enrichment procedure was performed using a 4 × 50 mm ProPac IMAC-10 analytical column (Thermo Fisher Scientific). For the JY cells an equivalent of 5 mg of protein was used for one IMAC purification while for the *C. difficile* cells, the total amount of peptides from each sample were applied, with an average of 11 (± 1.4 (S.D.)) mg protein per sample. Prior to each IMAC run the column was stripped using 50 mM EDTA/0.5 M NaCl pH 4 and charged with 25mM FeCl_3_ in 100mM acetic acid, according to the manufacturer’s instructions, using an offline pump system (Shimadzu). After this, the column was connected to an Agilent 1200 chromatography system running at a flow rate of 0.3 mL/min. Prior to sample loading, the column was equilibrated in solvent A (water/acetonitrile/trifluoroacetic acid 70/30/0.07 (v/v/v)) for 30 mins at a constant flow rate of 0.3 ml/min.

Tryptic peptides (resuspended in 260 µL solvent A, of which 250 µL injected) were loaded and non-phosphopeptides were removed by washing the column for 20 min at with solvent A. Peptide separation was performed using a linear gradient from 0 to 45% of solvent B (0.5% v/v NH_4_OH). The peak fraction of phosphopeptides between 46-49 min was manually collected and lyophilized prior to mass spectrometric analysis.

### LC-MS/MS analysis

Lyophilized peptides were reconstituted in 100 μL water/formic acid 100/0.1 (v/v) and analysed by on-line C18 nanoHPLC MS/MS with a system consisting of an Easy nLC 1200 gradient HPLC system (Thermo Fisher Scientific, Bremen, Germany), and an Orbitrap Fusion™ Lumos™ Tribrid™ mass spectrometer (Thermo Scientific™). Samples (10 μL, in duplicate) were injected onto a homemade pre-column (100 μm × 15 mm; Reprosil-Pur C18-AQ 3 μm, Dr. Maisch, Ammerbuch, Germany) and eluted on a homemade analytical nano-HPLC column (30 cm × 75 μm; Reprosil-Pur C18-AQ 3 um). The gradient was run from 2% to 36% solvent B (water/acetonitrile/formic acid 20/80/0.1 (v/v/v)) in 120 min. The nano-HPLC column was drawn to a tip of ∼5 μm which acted as the electrospray needle of the MS source.

The Lumos mass spectrometer was operated in data-dependent MS/MS mode for a cycle time of 3 seconds, with a HCD collision energy at 32 V and recording of the MS2 spectrum in the Orbitrap. In the master scan (MS1) the resolution was 120000, the scan range *m/z* 300-1500, at an AGC target of 400000 at a maximum fill time of 50 ms. Dynamic exclusion was set after n=1 with an exclusion duration of 60 s. Charge states 2-4 were included. For MS2, precursors were isolated with the quadrupole with an isolation width of 1.2 Da. First mass was set to 110 Da. The MS2 scan resolution was 30000 with an AGC target of 50000 at maximum fill time of 60 ms.

### Data analysis

MaxQuant software (version 1.5.1.2) was used to process the raw data files, which were searched against the *C. difficile* strain 630 database (4103 entries) or human canonical database (67911 entries). The mass tolerance for MS1 and MS2 was set to 4.5 and 20 ppm, respectively. Trypsin was selected as the enzyme with a maximum of two missed cleavages. Carbamidomethylation of cysteines was selected as a fixed modification and phosphorylation on serine, threonine and tyrosine, as well as oxidation of methionine and acetylation of the protein N-terminus were selected as variable modifications. The match between runs option was selected with a window of 2 min. The false discovery rate at the peptide level was set to 1% and for modified peptides a score cut-off of 40 was used.

The MaxQuant output table “phospho (STY)Sites.txt” (**Supplemental Table 1**) was used for further analysis of the phosphopeptides. For the stringent phosphorylation site assignment, only peptides with a localisation probability score >0.95 were selected. Because we used the match between runs option, we only used unique peptides without considering the phosphorylation site (i.e. not considering phospho-isomers) for comparison of different samples.

Gene Ontology protein/gene set enrichment analysis was performed using a sorted list of all identified phosphoproteins in each analysis time point, i.e. mid-exponential, beginning stationary, and late stationary growth phase. The sorting was based on the abundance of the phosphoproteins identified and determined by the ion intensity as reported by the mass spectrometer in the output of MaxQuant (cut-off at score of 40 and false discovery rate at 1%). We considered the phosphoproteins that were reported at least once in any of the three replicates. Each protein identification was considered only once in the sorted list with a rank determined by the most intense results we have obtained in any of the three replicates by any protein-associated phosphopeptide. The enrichment was performed using weighted Kolmogorov-Smirnov-like statistic as implemented in the common gene set enrichment analysis. We limited our analysis to the top 30 enriched Gene Ontology terms. We generated enrichment maps that represent the relationship between the enriched terms as a network. The edges of an enrichment map correspond to the number of shared proteins between the associated terms. These maps give a higher-level overview of the enrichment analysis and allow identifying functional modules with ontology terms that are related to each other by the underlying protein set used. For the functional analysis we used all Gene Ontology annotation of *C. difficile* as reported by the Gene Ontology database hosted by EBI (as of August 15 2021). For the functional analysis and visualization, we used R 3.6.2 and Bioconductor library Cluster Profiler.

### Data availability

All data are contained within the manuscript or the associated Supplementary Material (available via the journal website), or are available from the authors on request. The mass spectrometry proteomics data have been deposited to the ProteomeXchange Consortium via the PRIDE partner repository (37) with the dataset identifier PXD029475.

## RESULTS

### A one-step IMAC procedure allows the identification of phosphopeptides from *C. difficile*

The number of phosphorylated proteins in bacteria is generally much lower than in eukaryotic cells (1). To benchmark our IMAC workflow, we therefore first analysed a human cell line (JY cells) where we were expecting a relatively high number of phosphopeptides compared to *C. difficile*. From three biological replicates, we were able to identify a total of 10620 unique phosphopeptides with a good overlap between the different sample (**Supplemental Figure 2**). These results are in line with other recent studies (38, 39), and show that our approach allows the reliable identification of phosphopeptides in complex samples.

Next, we analysed the phosphoproteome of *C. difficile* at three different time points (mid-exponential phase, start stationary phase and late stationary phase (24 hr)) from three independently grown cultures. Overall, this resulted in the identification of almost 3000 phosphopeptides (**Supplemental Table 1**). Of these, 1759 had a high localisation probability (LP; above 0.95) for the correct assignment of the phosphorylation site. These sites were found on 1604 unique tryptic peptides derived from more than 700 proteins. Within this set of peptides, 75% was found to be phosphorylated on a serine, 20% on a threonine residues and 5% on a tyrosine (**Figure 1A**). Based on these results, we conclude that serine phosphorylation is most common in *C. difficile*, as also observed for other species (4, 11).

**Figure 1.**
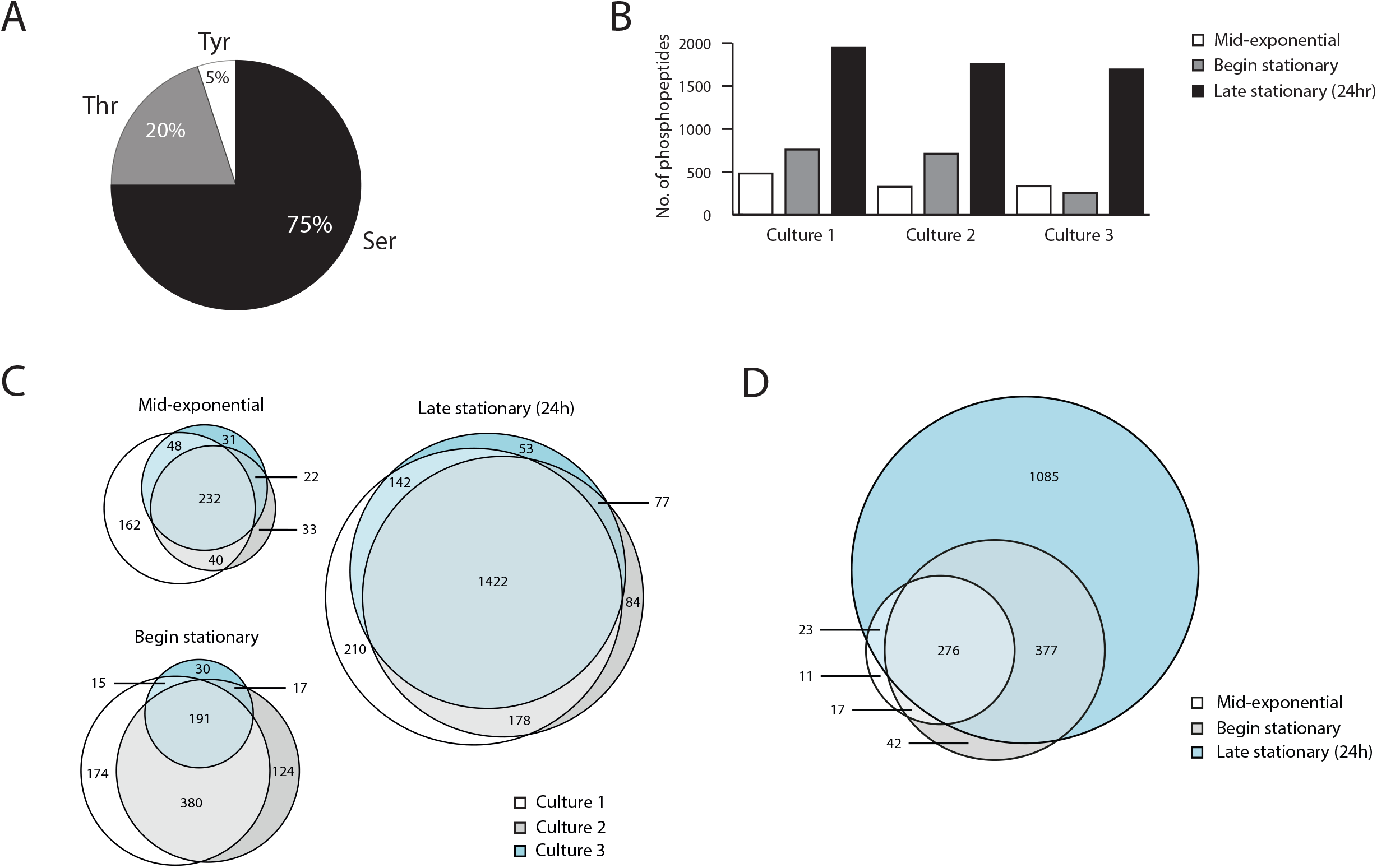
Global overview of phosphoproteins in *C. difficile*. **A**. Pie chart indicating the percentages of Ser, Thr and Tyr phosphorylation. **B**. Number of identified phosphopeptides over time for the three biological replicates analysed. For the definition of timepoints, see Supplemental Figure 1. **C**. Concordance between the phosphopeptide identifications between the three biological replicates. **D**. Changes in the identified phosphoproteome over time. Overlap in the identified phosphopeptides per timepoint for culture 2 is indicated in a Venn diagram.

To compare the different samples, we selected unique peptide sequences that were found to be phosphorylated, without considering phospho-isomers (i.e. phosphorylation at different sites in the same peptide). In general, we observed a progressive increase in protein phosphorylation during growth. We identified most phosphopeptides at later growth phases with the most prominent increase between the samples taken at the onset of stationary growth phase and the samples from the 24 hr cultures (**Figure 1B**).

We compared the reproducibility of the identifications between the different samples at the various timepoints. At the mid-exponential growth phase 232 phosphopeptides were identified in all three samples, corresponding to 41% of the total number of peptides identified at this timepoint (**Figure 1C**). For the second time point, 191 phosphopeptides were found in all three samples (20% of the total number identified at this time point). This relatively low percentage was mainly due to one of the samples, in which we found a lower number of phosphopeptides (**Figure 1B**, Culture 3) as the overlap between phosphopeptides was much higher between the other two cultures at this time point (63% of the total number). Of note, this was not due to experimental variation as analysis of a second sample from the same culture at the same time point showed a similar low number of phosphopeptides. After 24h, 1422 phosphopeptides were common to all three samples (66% of the total number identified at this time point) Overall, this shows significant congruence between our phosphopeptide identifications, despite individual processing of all three biological replicates and timepoints.

Within one culture, most phosphopeptides observed at the earlier time points were also observed at later stages (as exemplified for Culture 2 in **Figure 1D**), suggesting a gradual increase in the repertoire of phosphoproteins rather than a large scale reprogramming. This is also clear from the fact that 155 phosphopeptides were found in all samples at all time points. Nevertheless, we also observed a limited set of proteins that appear to be phosphorylated predominantly, if not exclusively, in the exponential growth phase (**Supplemental Table 1**) and are absent from late stationary growth phase samples. One such example is discussed further below.

We performed protein/gene set enrichment analysis of the phosphorylated proteins identified and determined the molecular functions and biological processes associated with these proteins. **Figure 2** represents the enrichment maps for each of the three time points, providing an overview by aggregating the enriched terms according to the shared protein set. Comparing the enrichment maps side by side revealed that during the first two stages there were concentrated, interconnected clusters of functions/processes, i.e. performed by same set of proteins; a trend that was lost in the late stationary growth phase. At the mid-exponential growth phase, these clusters are largely associated with metabolic and biosynthesis processes including few related to phosphorylation, as well as translation-associated functions and processes. At the beginning of the stationary growth phase, additional functions/processes start to be visible, including those related to ion/cation channel activities, ATP related processes, as well as cell motility and carbohydrate metabolic processes. At the late stationary growth phase, multiple parallel-unconnected functions and processes like DNA repair, transferases, catalytic activities were enriched, but also nucleotide biosynthetic processes that were already visible at the earlier time points. Of note, only the top 30 terms are shown in these maps, i.e. the functions and processes that are visible on top in the early stages likely are still active at the late stage, but they do not appear in the top 30.

**Figure 2.**
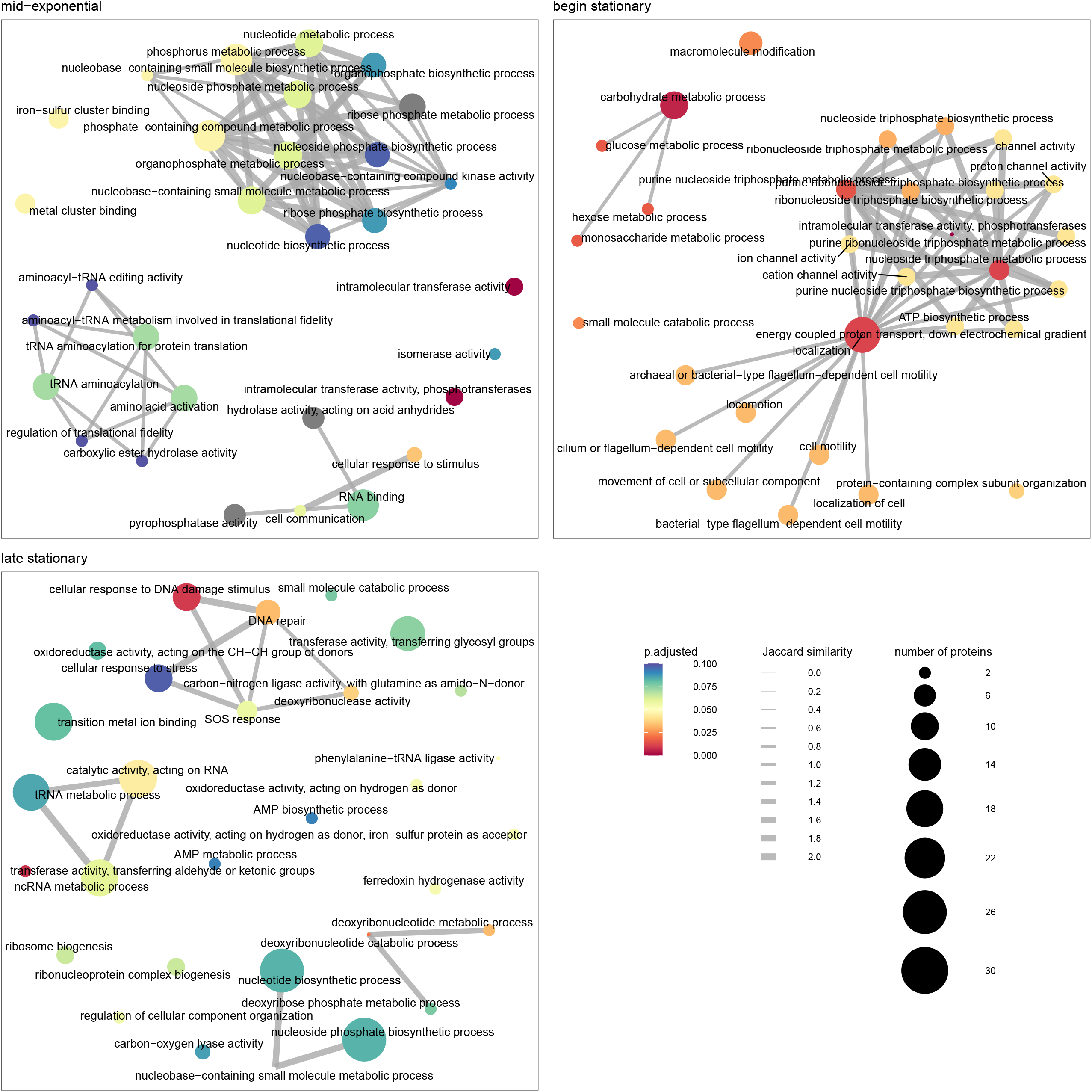
Enrichment maps of Gene Ontology terms obtained from *C. difficile* phosphoprotein set enrichment analysis. Three enrichment maps correspond to the three time points analyzed in this study (mid-exponential, beginning stationary, and late stationary growth phase); each map is limited to the top 30 enriched terms. The leaves represent Gene Ontology terms and the edges correspond to the Jaccard similarity between the leaves based on the shared proteins.

### Phosphorylation related to annotated kinases

We used the UniProt annotation of the *C. difficile* 630 genome (search with “cd630+kinase” in June 2019, 182 hits) to identify potential kinases and evaluate auto-phosphorylation or phosphorylation of substrates of these kinases on the basis of findings in other organisms. Here, we focus on the putative protein kinases that are expected to phosphorylate serine, threonine and tyrosine residues: SpoIIAB (CD630_07710)(32), RsbW (CD630_00100)(31), HPr kinase (CD630_34090)(30), PrkC/Stk (CD630_25780)(33) and CD630_21480. Other kinases identified by their annotation phosphorylate small molecules (nucleotides, metabolites), or are part of two-component regulatory systems and fall outside the scope of the present study. SpoIIAB is an anti-sigma-factor which in other bacteria is known to phosphorylate the anti-sigma factor F antagonist SpoIIAA during the process of sporulation (26, 32). In line with a role during sporulation, SpoIIAA phosphopeptides were exclusively identified in the 24 hr sample, but not in samples taken at mid-exponential or early stationary growth phase. Phosphorylation of SpoIIAA was found on Ser56 (LP=0.83), Ser57 (LP=0.55), Ser77 (LP=1.00), Ser82 (LP=0.84), Ser83 (LP=0.97) (**Supplemental Table 1**). Of these, Ser56 corresponds to the equivalent serine of *B. subtilis* SpoIIAA (Ser58) which is known to be phosphorylated by SpoIIAB in that organism (40). The corresponding peptide was found in all *C. difficile* cultures after 24 hrs. Manual inspection of the MS/MS spectrum **(Figure 3A)** confirmed phosphorylation of Ser-56 (corresponding to Ser14 in the tryptic peptide), because we observed neutral loss of phosphoric acid starting from the y_10._

**Figure 3.**
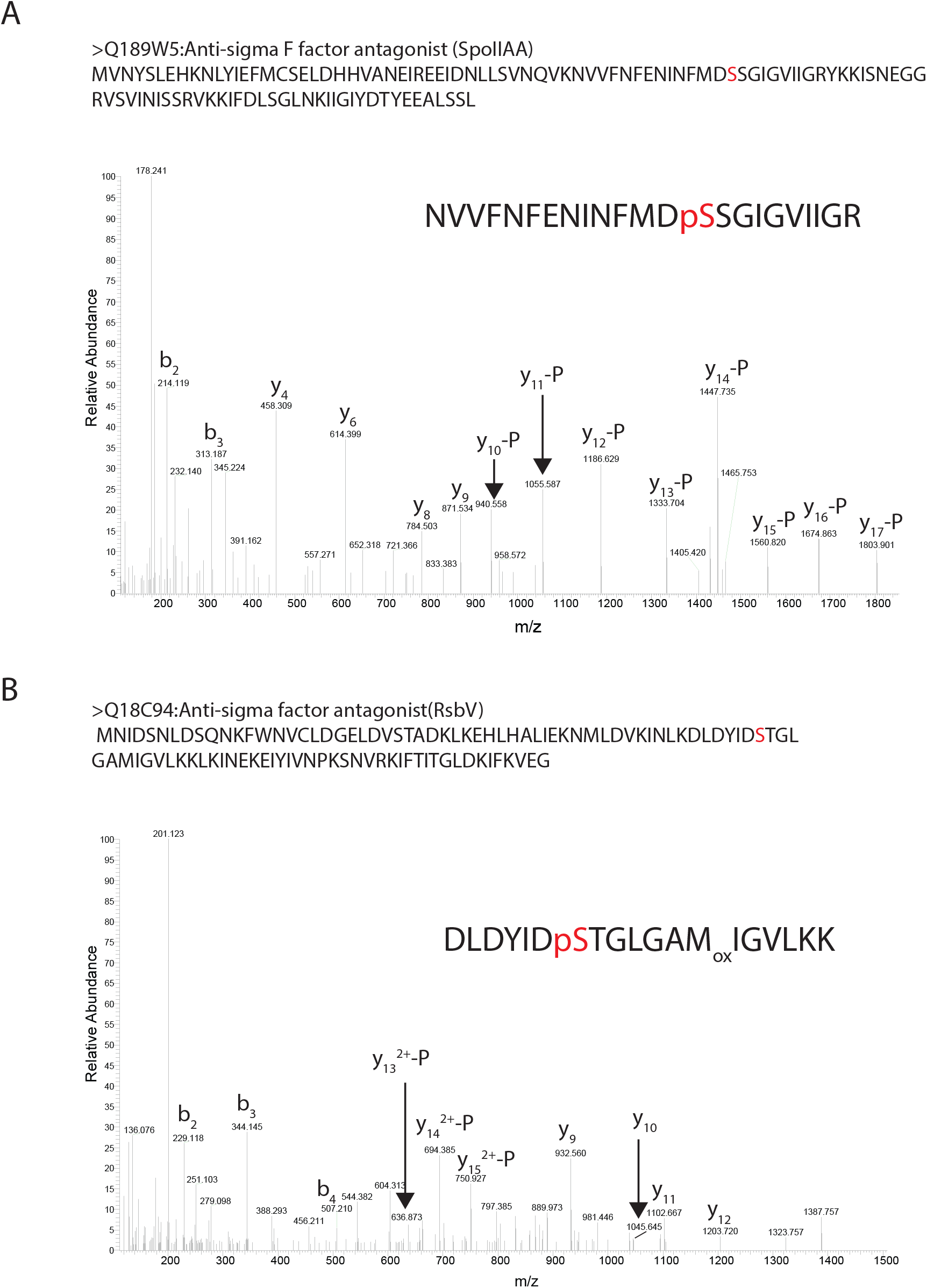
Spectra of conserved phosphorylation in regulatory proteins of *C. difficile*. **A**. MS/MS spectrum of the tryptic phosphopeptide NVVFNFENINFMDpSSGIGVIIGR (pS=phosphoserine) from SpoIIAA, demonstrating phosphorylation of Ser-14 (Ser-56 in the full length protein, indicated in red) **B**. MS/MS spectrum of the tryptic phosphopeptide DLDYIDpSTGLGAMIGVLKK (pS=phosphoserine) from RsbV, demonstrating phosphorylation of Ser-7 (Ser-57 in the full length protein, indicated in red). Ions assigned as “-P” have lost H_3_PO_4_

We also found phosphorylation of the kinase SpoIIAB itself (**Supplemental Table 1**): Ser13 (LP=0.98), Ser17 (LP=1.0) and Ser106 (LP=1.00). Lower confidence sites in SpoIIAB include Ser123 (LP=0.50), Ser124 (LP=0.49) and Thr111 (LP=0.62). Of all these sites, only Ser106 was found consistently in all three 24hr cultures and the assignment of the site was confirmed by manual inspection of the MS/MS data (**Supplemental Figure 3A)**.

RsbW is the anti-sigma factor for σ^B^, an important regulator of the stress response in gram-positive bacteria (27, 31). The interaction of RsbW with σ^B^ is inhibited through the binding of the anti-σ-factor antagonist RsbV, to RsbW (27, 31). The interaction between RsbV and RsbW is negative regulated by RsbW-dependent phosphorylation of RsbV (27). We found two high-LP phosphopeptides for RsbV, indicative of phosphorylation of Ser57 (LP=0.93) and Ser84 (LP=1.00) and a low-LP phosphorylation of Thr58 (LP=0.56). Work in *B. subtilis* has identified three phosphorylation events and two of these (Ser 56 and Thr57) are on the residues equivalent to the ones identified here in *C. difficile* RsbV (15). On the basis of manual inspection of the MS/MS data, we provide strong evidence for phosphorylation of Ser57 in *C. difficile* RsbV **(Figure 3B)**. We also identified a significant number of phosphopeptides derived from the kinase RsbW itself, in most cultures, at the 24h timepoint (**Supplemental Table 1**): Ser87 (LP=0.87), Ser89 (LP=0.83), Thr90 (LP=0.79) and Ser 99 (LP=1.00).

HPr kinase (HprK/ScoC) can phosphorylate the phosphocarrier protein HPr (PtsH), a protein involved in the import of monosaccharides as part of the PTS system (5). In *B. subtilis*, HPr kinase-mediated phosphorylation of HPr occurs on Ser46 (41, 42). We found phosphorylation of the homologous residue (Ser45, LP=1.00) of *C. difficile* HPr in all samples, and additionally identified high confidence phosphorylation of Ser11 (LP=1.00), and lower confidence phosphorylation of Ser30 (LP=0.85) and Thr31 (LP=0.82) (**Supplemental Table 1**). The equivalent site of Ser11 in *B. subtilis* HPr (Ser12) was also identified in a phosphoproteomics experiment (15). Importantly, the corresponding tryptic peptide of *C. difficile* HPr also contains a conserved histidine (His14), which is known to be phosphorylated through the phosphotransferase activity of the PTS-system (5, 43). However, the MS/MS spectrum of the tryptic peptide clearly demonstrates neutral losses of phosphoric acid from the b_3_-ion and a series of non-phosphorylated y-ions until y_12_. Based on this, and the absence of the characteristic pHis immonium ion in the MS/Ms spectrum, we conclude that Ser12 rather than His14 of HPr is phosphoryalted (**Supplemental Figure 3B**). We also found a single phosphopeptide derived from the HprK kinase itself: Thr299 (LP=1.00) appears to be phosphorylated primarily in stationary growth phase, with most peptides identified after 24h.

Phosphorylation of the Hanks-type serine-threonine kinases *C. difficile* PrkC (33) and CD630_21480 is hitherto undescribed and their substrates are virtually uncharacterized. We did not identify phosphopeptides derived from CD630_2148, but found extensive phosphorylation of PrkC in the region between Ala158 and Lys183 (tryptic peptide AV**<u>S</u>**N**<u>ST</u>**M**<u>T</u>**NIG**<u>S</u>**IIG**<u>S</u>**VHYFSPEQAK, LP>0.95 in **<u>red</u>** and 0.70<LP<0.95 in **<u>bold</u>**) throughout growth. Notably, the conserved residues Thr163 and Thr165 were also found to be phosphorylated in *B. subtilis* (44). Thr290 of *B. subtilis* PrkC, another residue that was identified in multiple studies to be phosphorylated (15, 44), is not conserved in *C. difficile*. However, two residues in the equivalent region of *C. difficile* PrkC (Thr287 and Thr302, both LP=1.00) were identified, suggesting that phosphorylation in this region is structurally conserved. We also found phosphopeptides that indicate phosphorylation close to the N-terminus of PrkC (Thr4 and Thr35, LP=1.00) (**Supplemental Table 1**).

A single substrate for PrkC has been characterized: CwlA (CD1135) (34). For this protein, phosphorylation was reported on Ser136 and Thr405, with the latter being specific for PrkC. In our analyses, we find multiple Ser-phosphorylated peptides, including Ser136 (LP=1.00), and Thr405 (LP=1.00) (**Supplemental Table 1**). Additionally, it appears that PrkC is involved in cell wall homeostasis and antimicrobial resistance based on phenotypes of a *prkC* knockout strain (33). The authors postulate that part of the effects could be mediated by CD630_12830, a homolog of IreB of *Enterococcus faecalis*. In this organism, IreB negatively controls cephalosporin resistance and is phosphorylated by the serine/threonine kinase IreK on a conserved N-terminal threonine residue (45). We confirmed phosphorylation of the equivalent residue (Thr8, LP=1.00) across all time points, in all samples (**Supplemental Table 1**).

### Dynamic phosphorylation within a protein

PrkC may also regulate cell division, such as DivIVA (33, 46-49). We identified high-confidence phosphopeptides derived from *C. difficile* DivIVA (CD630_26190), indicating phosphorylation of Ser75 (LP=1.00), Ser82 (LP=1.00) and Thr160 (LP=0.98). The phosphorylation of DivIVA is one of the few examples of consistent phosphorylation at a specific site (Thr160) at early time points, but not after 24 hr of culturing (**Supplemental Table 1**). Interestingly, the cell division protein SepF (CD630_26220), which is encoded in the same gene cluster as *divIVA*, shows the same pattern of phosphorylation for Thr56 (LP=0.96). In contrast, the other sites in DivIVA (Ser75 and Ser82) were found to be consistently phosphorylated in stationary growth phase, but never at the mid-exponential phase. It therefore appears that the pattern of phosphorylation of DivIVA differs throughout growth. Exponential growth phase phosphorylation is not observed for all cell division proteins, as for instance ZapA (CD630_07910) and FtsZ (CD630_26460) are predominantly phosphorylated in stationary growth phase (**Supplemental Table 1**).

We were interested to see if more proteins demonstrate dynamic phosphorylation. Manual inspection of the list of phosphopeptides identified two hypothetical proteins that show different patterns of phosphorylation in exponential compared to stationary growth phase: CD630_27170 and CD630_28470 (**Supplemental Table 1**). Phosphorylation of CD630_27170 occurs exclusively on Ser573 (LP=1.00) in exponential growth phase, at Ser573 and Ser286 (LP=1.00) at the onset of stationary growth phase, and at multiple other sites at 24h. CD630_28470 is phosphorylated at Ser186 (LP=1.00) in exponential growth phase and onset of stationary phase but not at late stationary phase (**Supplemental Table 1**). Together, these examples indicate that post-translational modification within a particular protein in *C. difficile* can change in a growth phase dependent manner.

### Indirect detection of cysteine phosphorylation

In addition to protein kinase-dependent phosphorylation, in which the phosphate group is donated by ATP, bacteria can also use the phosphate group from phosphoenolpyruvate (PEP) in a process catalysed by a phosphotransferase system (PTS). This is of particular relevance for the import of monosaccharides into the cell, during which the sugars are converted to phospho-sugars (5). The initial enzymes involved in this cascade of events (Enzyme I (PtsI) and Hpr (PtsH) are phosphorylated at conserved histidine residues and do not have specificity for the different monosaccharides. The subunits of the Enzyme II complex (EIIA, EIIB and EIIC) provide the necessary specificity for different monosaccharides. In our phosphoproteomic dataset, we found that many of these subunits of the different EII complexes are phosphorylated at serine and threonine residues (**Supplemental Table 1**). Activation of the mono-saccharide during uptake, however, requires the transfer of phosphate from a conserved cysteine residue of the EIIA and EIIB subunits, respectively. We accidently put carbamidomethylation of Cys as a variable, instead of a fixed, modification during one of our database searches and serendipitously observed cysteine-containing peptides from EII complex subunits that did not appear to be phosphorylated (as expected based on our enrichment strategy) or carbamidomethylated (as expected from our proteomics workflow). Of note, when we searched data from a HeLa tryptic digest, that we use as a standard for our LC-MS/MS setup, with the same parameters, we find very few free cysteines (18 out of 2753 peptide spectrum matches of peptides containing a cysteine).

As an example of a tryptic peptide with a free cysteine in our *C. difficile* data, we observed the tryptic peptide ILVA**<u>C</u>**GAGIATSTIV**<u>C</u>**DRVER from the PTS system IIB component (CD630_10830, aa 4-24) containing two cysteines (in bold and underlined). In total more than 300 peptide spectral matches for this peptide were identified in all our LC-MS/MS analyses of *C. difficile* phospho-samples. In all these identifications, the second cysteine (aa 16 in the tryptic peptide) was found to be carbamidomethylated, while the first cysteine at position 5 was a free cysteine **(Supplemental Figure 4**). Importantly, the first cysteine corresponds to the active site cysteine that is transiently phosphorylated in the P-loop of EII subunits (50). We therefore hypothesize that the identification of such a peptide is due to the fact that these cysteines were phosphorylated during the initial steps of our workflow. This would allow for enrichment during the IMAC procedure and – if the phosphorylation persists during the reduction and alkylation steps, preclude carbamidomethylation. As no phosphorylated cysteine modifications were found in the LC/MS-MS, it appears that the phosphate group is lost during the later stages of our workflow (due to instability of the phosphorylated cysteine under our experimental conditions).

In conclusion, it appears that a one-step Fe-IMAC enrichment procedure can indirectly detect at least a subset of cysteine-phosphorylated proteins.

## DISCUSSION

In this study we characterized the phosphoproteome of a laboratory strain of *C. difficile* at three different timepoints. We show that many proteins of *C. difficile* are phosphorylated in stationary growth phase, and an analysis of subset of the identified phosphoproteins indicates that at least some of the phosphorylation sites are conserved between phylogenetically distinct organisms.

Many different methods have been used to analyse bacterial phosphoproteomes (1, 2, 11). A crucial step in these analyses is the method used to enrich for phosphopeptides (2, 4). In our study, we employed Fe-IMAC as this has recently been shown to be a very effective way for analyzing bacterial phosphoproteomes (38, 39). The number of phosphoproteins identified here is in line with these studies, and indicates this is a suitable method for the analysis of phosphoproteins in *C. difficile*. We identified Ser-, Thr-, and Tyr-phosphorylation and provided indirect evidence for Cys-phosphorylation (**Figure 1A, Supplemental Table 1**). Our workflow does not allow the identification of His-phosphorylation, but recent studies have shown that the Fe-IMAC method can be adapted to detect the thermodynamically less stable His-phosphorylation for future experiments (51, 52). The identification of other forms of phosphorylation, such as Arg phosphorylation, requires different approaches (53). We also observed multiply phosphorylated proteins (**Supplemental Table 1**), as has seen in other organisms as well (12, 54-56).

Our work provides a starting point for experimental validation of the identified phosphoproteins, as well as a dissection of the contribution of individual protein kinases and phosphatases. Generally, the action of protein kinases is better understood than that of phosphatases (16). We identified site-specific phosphorylation in the protein kinase PrkC (**Supplemental Table 1**). A knockout of the gene encoding this kinase demonstrates pleiotropic effects (33), and comparing wild type and *prkC* negative cells using our approach may help to elucidate the substrates of this protein.

The identified phosphopeptides also offer a possibility of characterizing the effect of phosphorylation on substrate proteins, by mutating residues in these proteins to non-phosphorylatable analogs (alanine for serine/threonine and phenylalanine for tyrosine) or phosphomimetic negatively charged amino acids (aspartate and glutamate) (2, 10). Such an approach may be preferable over mutating the kinases and phosphatases, as these can have overlapping specificities (57).

In previous phosphoproteomic analyses, a large divergence was often observed in the lists of phosphorylated proteins even within a single genus or species (2, 58, 59). In our experiments, we observed apparently conserved phosphorylation events on several proteins, including SpoIIA, SpoIIAB, RsbV and others (**Figure 3, Supplemental Table 1**), which may appear in contrast with these observations. Part of the reason for the limited overlap in previous experiments may have been the diversity in enrichment methods and experimental protocols, limited sensitivity of the workflows and variability in sampling method and timepoint. Our workflow allows the reliable identification of a large set of phosphoproteins in comparison with other studies (11, 12, 16, 17). Moreover, we applied rigorous quality control of phosphosite assignment. It is likely that developments in both mass spectrometry instrumentation, data acquisition and analysis pipelines will reveal that many more processes in fact may be regulated by conserved phosphorylation events. For example, a recent phosphoproteomic analysis of *Staphylococcus aureus* increased the number of phosphosite identifications over 20-fold compared to earlier analyses, thereby also increasing the number of processes affected (38). Cross-species comparisons may be facilitated by dedicated databases of phosphorylated proteins, such as PHOSIDA and dbPSP 2.0 (60, 61) and ultimately lead to improved prediction of phosphorylation in bacteria (62).

Our work indicates that protein phosphorylation in *C. difficile* may differ both spatially (within a protein) and temporally (between timepoints). Specifically, we observe low levels of phosphorylation in exponential growth phase, whereas phosphorylation increases markedly upon entry into stationary growth phase. Of note, we observed a highly variable number of phosphoproteins at the entry into logarithmic growth phase, with two out of three samples showing clearly elevated levels, whereas a single biological replicate resembled the profile observed in exponential growth phase (**Figure 1B**). This suggests that the phosphorylation state could change rapidly, and supports the notion that sampling timepoint could explain differences between phosphoproteomic datasets. We also report that specific phosphorylation patterns can be observed within a protein, as exemplified by DivIVA and SepF (**Supplemental Table 1**). This has also been observed for other organisms; for instance, in *Streptomyces coelicolor*, 58/85 phosphorylation sites were differentially phosphorylated during differentiation (13). Interestingly, Thr160 of *C. difficile* DivIVA is located in the putative tetramerization region of the protein (63) and phosphorylation at the C-terminus of DivIVA has also been observed for *S. pneumoniae*, where is it mediated by the eSTK protein StkP (49). Mutations of this region lead to aberrant cell morphology, suggesting that phosphorylation in of Thr160 in *C. difficile* might have functional consequences. Thr56 of *C. difficile* SepF is directly adjacent to the predicted compact domain that mediates self-interaction and interaction with the cell division protein FtsZ (64). The functional consequences of the dynamic phosphorylation of DivIVA and SepF remain to be elucidated.

We focused our in-depth analyses on those proteins for which the phosphosite assignment could be done with high probability; where necessary, we manually verified the automatic identifications. Nevertheless, we also observed peptides for which it was not possible to assign the modification to a particular residue, similar to others (59). For those proteins, alternative fragmentation techniques (65) or processing of the sample – using for instance proteases different than trypsin – might yield better results.

The phosphoproteome described here complements existing omics approaches for *C. difficile*, such as genomics, transcriptomics, proteomics and metabolomics. Integration of these data may lead to a systems-level understanding of *C. difficile* physiology and – more broadly – may contribute to our understanding of the role of phosphorylation in the regulation of bacterial pathogenesis (16).

## Acknowledgements

The authors wish to acknowledge J. Corver for helpful discussions, and I. Martin-Verstraete for communicating results prior to publication.

## LIST OF ABBREVIATIONS

HPLC: High Performance Liquid Chromatography
LC: Liquid Chromatography
LC-MS: Liquid Chromatography – Mass Spectrometry
nLC-MS/MS: Nano Liquid Chromatography -Tandem Mass Spectrometry
LP: localisation probability
aa: amino acid

## LEGENDS

**Supplemental Figure 1. Overview of growth and sampling of the *C. difficile* cultures. A**. Growth curve indicating obtained optical densities at 600nm and sampling timepoints for this study. **B**. Table summarizing protein yield from 100 mL (time point 1) and 50 mL (time points 2 and 3) of sample.

**Supplemental Figure 2**. Summary of total phosphopeptides retrieved from JY cells. Overlap between the three biological replicates is represented with the Venn diagram.

**Supplemental Figure 3. Spectra of conserved phosphorylation in regulatory proteins of *C. difficile***. A. MS/MS spectrum of the tryptic phosphopeptide AMEPLYTSKPELDRpSGMGFTVMK) from SpoIIAB, demonstrating phosphorylation of Ser-15 (Ser-106 in the full length protein, indicated in red) B. MS/MS spectrum of the tryptic phosphopeptide NApSGLHARPAGMFVK from HPr, demonstrating phosphorylation of Ser-3 (Ser-11 in the full length protein, indicated in red). pS=phosphoserine. Ions assigned as “-P” have lost H_3_PO_4_

**Supplemental Figure 4. Indirect identification of cysteine phosphorylation**

MS/MS spectrum of the tryptic peptide LVACGAGIATSTIVC_cam_DRVER from the phosphotransfer system (PTS) component IIB (CD630_10830) in which the first cysteine (in red) is a free cysteine while the second is carbamidomethylated (cam).

**Supplemental Table 1**. Full data set of the phosphoproteome analysis and site assignment for Ser, Thr and Tyr phosphorylation in *C. difficile*.

